# *Slc35a2* mosaic knockout impacts cortical development, dendritic arborisation, and neuronal firing in the developing brain

**DOI:** 10.1101/2024.02.18.580216

**Authors:** James Spyrou, Khaing Phyu Aung, Hannah Vanyai, Richard J Leventer, Snezana Maljevic, Paul J Lockhart, Katherine B Howell, Christopher A Reid

## Abstract

**Objective:** Mild malformation of cortical development with oligodendroglial hyperplasia in epilepsy (MOGHE) is an important cause of drug-resistant epilepsy. A significant subset of individuals diagnosed with MOGHE display somatic mosaicism for loss-of-function variants in *SLC35A2*, which encodes the UDP-galactose transporter. We developed a mouse model to investigate the mechanism by which disruption of this transporter leads to a malformation of cortical development.

**Methods:** We used *in utero* electroporation and CRISPR/Cas9 to knockout *Slc35a2* in a subset of layer 2/3 cortical neuronal progenitors in the developing brains of fetal mice to model mosaic expression.

**Results:** Histology of brain tissue in the mosaic *Slc35a2* knockout mice revealed the presence of upper layer-derived cortical neurons in the white matter. In contrast, oligodendrocyte patterning was unchanged. Reconstruction of single filled neurons identified altered dendritic arborisation with *Slc35a2* knockout neurons having increased complexity. Whole-cell electrophysiological recordings revealed that *Slc35a2* knockout neurons display reduced action potential firing and increased afterhyperpolarisation duration compared with control neurons. Mosaic *Slc35a2* knockout mice also exhibited significantly increased epileptiform spiking and increased locomotion.

**Interpretation:** We successfully generated a mouse model of mosaic *Slc35a2* deficiency, which recapitulates features of the human phenotype, including impaired neuronal migration. We show that knockout in layer 2/3 cortical neuron progenitors is sufficient to disrupt neuronal excitability and increase epileptiform activity and hyperactivity in mosaic mice. Our mouse model provides a unique opportunity to investigate the disease mechanism(s) that underpin MOGHE and facilitate the development of precision therapies.

## 1. INTRODUCTION

Mild malformation of cortical development with oligodendroglial hyperplasia in epilepsy (MOGHE) is a malformation of cortical development (MCD), which causes drug-resistant epilepsy with developmental delay and intellectual disability^1^. Epileptic spasms are the most common seizure type, with onset typically in infancy or early childhood^1, 2^. The epilepsy in individuals with MOGHE is often that of a developmental and epileptic encephalopathy, where uncontrolled epilepsy progressively impacts development^3^. As with many other MCDs, resection of the affected tissue may be required to successfully treat seizures and halt developmental decline.

MOGHE displays a number of histopathological hallmarks, including significantly increased numbers of heterotopic neurons in the white matter and along the grey-white matter boundary. Clusters of oligodendroglial cells are observed in both the white matter and deep layers of the cortex, in addition to areas of patchy myelination and a blurred grey-white matter junction^1, 2^. Somatic pathogenic variants in the X-linked gene *SLC35A2* were recently identified in brain tissue resected from a cohort of paediatric patients with MOGHE who underwent surgery as treatment^2^. Protein-truncating *SLC35A2* variants account for the majority of pathogenic variants identified in patients with MOGHE^2^. The mechanism by which loss-of-function *SLC35A2* variants and the resultant neuronal and oligodendroglial cortical abnormalities generate seizures remains unclear.

*SLC35A2* encodes the uridine diphosphate (UDP)-galactose transporter, which is responsible for the transport of galactose from the cytosol into the lumen of the Golgi apparatus and has a fundamental role in the glycosylation of proteins and lipids^4, 5^ (Figure 1A). Within affected cells, proteins and sphingolipids produced have truncated glycan chains due to the lack of galactose availability in the Golgi apparatus^4–6^. Glycosylation, especially N-linked glycosylation, is involved in a wide array of critical biological processes. These include protein folding, stability, and localisation^7^, as well as membrane excitability, cell migration and adhesion in the central nervous system specifically^8–10^.

**Figure 1.**
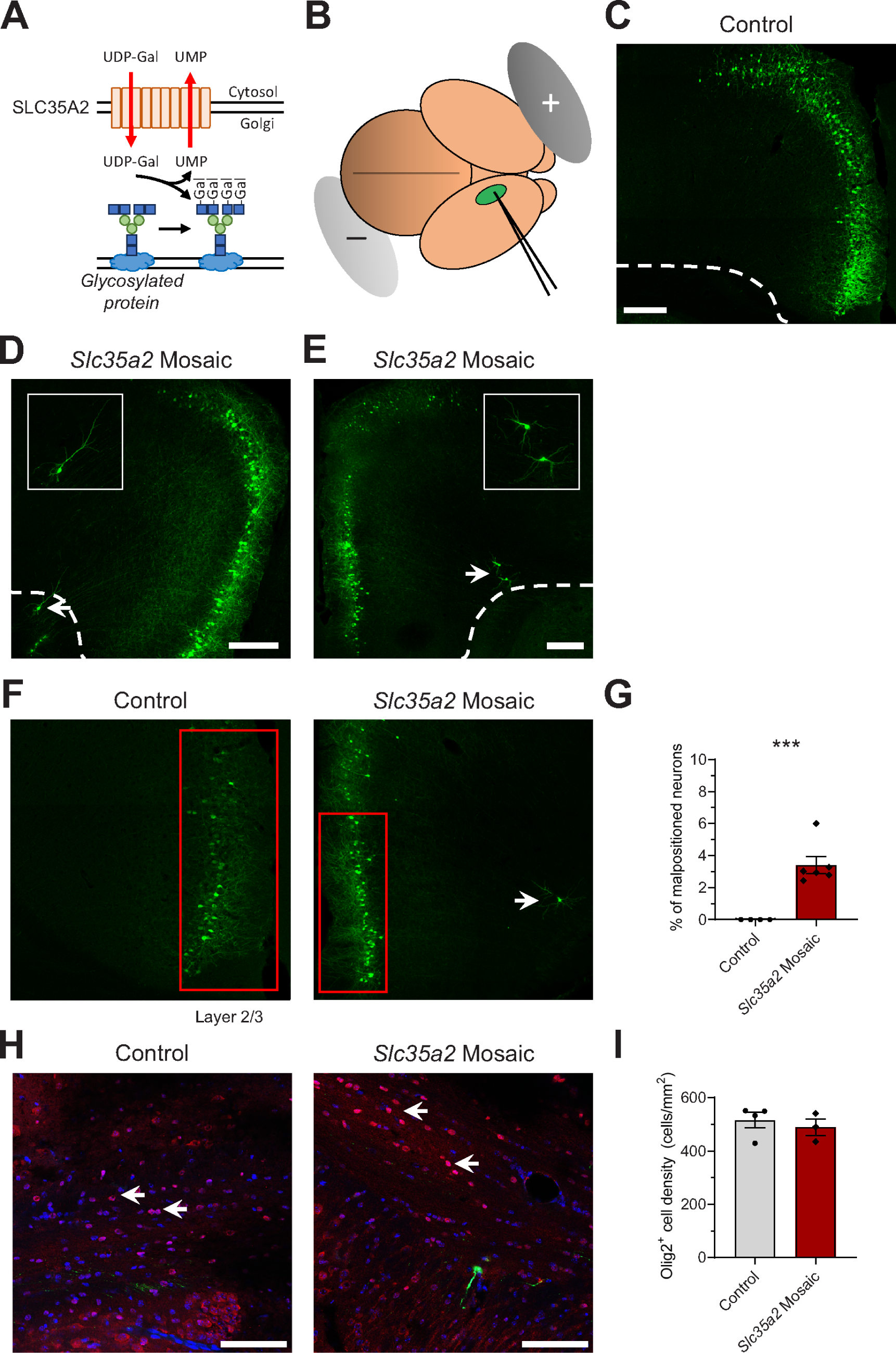
Mosaic *Slc35a2* KO mice display heterotopic neurons in the cortex and white matter. **(A)** Diagram of the function of the SLC35A2 protein in transporting UDP-galactose (UDP-Gal) across the membrane of the Golgi apparatus from the cytosol, where galactose is then attached onto growing glycan chains in the process of glycosylation. UDP = uridine diphosphate, UMP = uridine monophosphate. **(B)** Diagram of the placement of electrodes, and location of microinjection (into one of the lateral ventricles), for an E15±0.5 fetal mouse brain as viewed from above, facing right. **(C-E)** Representative confocal images of GFP-expressing neurons in brains of control **(C)** and mosaic *Slc35a2* KO **(D, E)** electroporated mice. Arrows indicate heterotopic neurons and dotted lines approximate grey-white matter boundaries. Insets: magnified images of heterotopic neurons. Scale bars = 250 µm. Mice underwent transcardial perfusion at ages P28-42. **(F)** Neurons located deeper than layer 2/3 when beyond 300 µm from the pial surface (boxes; left panel = control mouse, right panel = mosaic *Slc35a2* KO mouse; arrow indicates misplaced neuron). **(G)** Proportion of misplaced GFP-expressing neurons in control and mosaic *Slc35a2* KO mice ***P < 0.001 **(H)** Representative confocal images of Olig2-positive (Olig2^+^) cells (red fluorescence; white arrows) in the white matter of a control (left) and a mosaic *Slc35a2* KO (right) mouse, showing GFP fluorescence of a heterotopic neuron (green) in the latter, co-stained with DAPI (blue). Scale bar = 100 µm. **(I)** Quantification of Olig2^+^ cell density in the white matter of control and mosaic *Slc35a2* KO mice, showing no significant difference between the two groups.

Here we utilised *in utero* electroporation (IUE) and CRISPR/Cas9 to selectively knockout *Slc35a2* in a subset of cortical neuronal progenitors in the developing brain, resulting in mice with mosaic loss-of-function for *Slc35a2* in layer 2/3 excitatory neurons. We demonstrate a critical role for *Slc35a2* in supporting correct neuronal migration, consistent with recent data from Elziny *et al.*^11^. We extend these findings by performing a combination of behavioural, histological, electrographic, and biophysical assays to investigate the impact of this mosaicism on seizure generation and neuron function, to provide insight into the pathogenesis of MOGHE.

## 2. MATERIALS AND METHODS

### 2.1 Mice

Mice were ordered from the Animal Resources Centre (ARC; WA, Australia). Swiss strain mice were used for all experiments; C57BL/6J mice were additionally used for electrocorticography experiments. Different background strains are known to have different seizure thresholds, with C57BL/6J mice typically showing more severe seizure phenotypes compared with other common strains^12^. Mice were housed in standard 15 × 30 × 12 cm cages in a 12 hour light and dark cycle, with access to dry pellet food and tap water *ad libitum*. For all experiments both male and female mice were used, with age-matched cohorts and littermates used wherever possible. All experiments were approved by the Animal Ethics Committee at the Florey Institute of Neuroscience and Mental Health, and are reported in compliance with the ARRIVE guidelines^13^.

### 2.2 In Utero Electroporation and Plasmids

The IUE and microinjection procedures described below were based on those previously used by Hsieh *et al.*^14, 15^ (Figure 1B). Time-mated pregnant mice (at embryonic day E15±0.5) were anaesthetised with isoflurane and surgically incised to expose fetuses. Approximately 1 µL of plasmid solution, detailed below, was injected into the lateral ventricle of each fetal mouse, which were then electroporated to allow plasmid uptake from the ventricle into layer 2/3 cortical neuronal progenitor cell populations in the medial prefrontal cortex (mPFC).

A CRISPR knockout kit for *Slc35a2* was obtained from OriGene. The plasmid in this kit (pCas-Guide-Slc35a2) encodes humanised Cas9 and a mouse *Slc35a2*-specific single guide RNA (sgRNA). Sanger sequencing was performed at the Australian Genome Research Facility (AGRF, Melbourne, VIC, Australia) to verify the sgRNA sequence as GGTTGGTGGATCTACCGCTG. The specificity for Mouse *Slc35a2* was validated by comparison with the NCBI Reference Sequence NM_078484.3. An additional plasmid encoding green fluorescent protein (GFP; pCAG-GFP; GenScript), was used to label successfully transfected cells. Injection solutions were composed as follows in nuclease-free water: 1.5 µg/µL pCas-Guide-Slc35a2, 0.5 µg/µL pCAG-GFP, and 0.1% (w/v) Fast Green Dye; control injection solution consisted of only pCAG-GFP and Fast Green Dye.

### 2.3 Brain Dissociation, Fluorescence-Activated Cell Sorting, and MiSeq

Brain dissociation and MiSeq were performed at the Walter Eliza Hall Institute (WEHI; Parkville, VIC, Australia). Electroporated mouse pups aged P0-2 were euthanised and GFP-positive (GFP^+^) brain regions were isolated using a fluorescence dissecting microscope. Three control brains, transfected with GFP only, and six pCas-Guide-Slc35a2 brains with GFP^+^ cells in the mPFC were collected and processed. One GFP-negative control brain was used as a negative control. Brain tissue was treated with Trypsin/EDTA (Merck) for 15 minutes and then triturated. Tissue was then passed through 70 µm MACS SmartStrainers, and Miltenyi Debris Removal Solution was added according to the manufacturer’s instructions. Tubes were centrifuged and the top layer and debris were aspirated. Cell pellets were resuspended, filtered into fluorescence-activated cell sorting (FACS) tubes, and centrifuged at 300 g for 10 min. The supernatant was then aspirated, and the pellet was resuspended in 0.5% (w/v) BSA in phosphate-buffered saline (PBS). Cell suspensions were sorted on a BD FACSAria III Cell Sorter (BD Biosciences) at the Melbourne Cytometry Platform (Parkville, VIC, Australia). The negative control and a GFP-only control were used to establish gating for GFP^+^ cells. Positive and negative fractions were then sorted and collected from each sample. Cells were pelleted and resuspended in Viagen DirectPCR Lysis Reagent (Mouse Tail) with Proteinase K. Samples were incubated in a heated shaker at 55°C overnight before inactivation for 45 min at 85°C. CRISPR/Cas9 editing efficacy was determined by targeted PCR of the Cas9-targeted region and secondary PCR using overhang sequences, followed by Illumina MiSeq sequencing, as previously described^16^. Unique amplicons per sample were included where 20 or more reads were recorded. Amplicons were aligned to reference genomes using the UCSC Genome Browser Blat tool (https://genome.ucsc.edu/cgi-bin/hgBlat) and assessed for indels around the predicted guide cut site.

### 2.4 Transcardial Perfusions, Immunohistochemistry, and Imaging

Transcardial perfusions and immunohistochemistry (IHC) were performed as previously described^17^. At age P28-42, a subset of electroporated mice underwent transcardial perfusion, and their brains were cryoprotected in a solution of 30% (w/v) sucrose. Coronal brain sections were sliced with a Leica CM1950 cryostat at a thickness of 30-40 µm. Slices were imaged using a Zeiss LSM 780 confocal microscope to visualise cells expressing GFP. A subset of slides underwent further analysis through IHC. After incubation with primary and secondary antibodies, ProLong™ Gold Antifade mounting medium (Thermo Fisher Scientific) was added. Confocal images were processed with ImageJ (https://imagej.nih.gov/ij/). Neuronal morphology was assessed with the SNT plugin for ImageJ^18^; morphological features were measured and Sholl analyses were performed using semi-automated traces of dendrites. Neuron soma size was measured by tracing the soma of randomly selected GFP^+^ cells. Neuron positioning (% cells in layer 2/3) was quantified by counting the number of GFP^+^ cells both within and beyond 300 µm from the pial surface. Cells within this area were considered as correctly located in layer 2/3 of the cortex^14, 19^ and those beyond were considered to be incorrectly placed; from this a proportion of correctly placed cells was calculated. Olig2-positive (Olig2^+^) cell density was calculated by counting the number of cells immunoreactive for Olig2 in the white matter in confocal images of mouse brain slices and dividing this number by the area of the white matter in each respective image in mm^2^. A complete list of antibodies and fluorescent probes used can be found in Table S1.

### 2.5 Electrocorticography

Electrocorticography (ECoG) experiments were performed as previously described^17, 20^. After electrode implantation surgery, mice were allowed to recover for at least one week prior to recording. Mice were recorded for at least 7.5 hours, covering both the light and dark phases of the light-dark cycle. Raw ECoG data were visually assessed to identify epileptiform spikes, and the average number of spikes per hour was calculated for each mouse.

### 2.6 Open Field Exploratory Locomotion Assay

Locomotion and thigmotaxis were measured as previously described^17, 20^. Prior to all behavioural experiments, mice were acclimatized for 1 hour in a dimly lit behavioural room.

### 2.7 Brain slice electrophysiology and dendritic visualisation

Slice electrophysiology was performed as previously described^17, 21^. Coronal slices were prepared from the mPFC of male and female Swiss mice (P20-33). Brains were extracted into ice-cold slicing solution, containing (in mM) 87 NaCl, 3 KCl, 0.25 CaCl_2_, 3 MgCl_2_, 25 NaHCO_3_, 1.25 NaH_2_PO_4_, 75 sucrose, and 25 glucose (305 mOs/kg). Slices were collected from each brain, which were transferred to a holding chamber containing artificial cerebrospinal fluid (aCSF) comprising (in mM) 125 NaCl, 3 KCl, 2 CaCl_2_, 1 MgCl_2_, 25 NaHCO_3_, 1.25 NaH_2_PO_4_, and 25 glucose (310 mOs/kg), supplemented with 3.6 mM pyruvate and 1.7 mM ascorbate.

Whole-cell recordings were made from pyramidal neurons located in layer 2/3 of the medial prefrontal cortex (mPFC), in current clamp mode to measure firing properties. Transfected cells were identified by visualising their GFP fluorescence using an Olympus BX51WI microscope (Olympus, Tokyo, Japan). The identified cell was then approached with a strong positive pressure (∼40 mbar) and the electrode resistance was confirmed to be in the range 5-8 MΩ. A family of 1000 ms-long current steps was then applied, with increments of 20 pA (a total of 30 steps, repeated every 5 sec). Patching of heterotopic neurons in the white matter was attempted but were not able to be reliably visualised due to the scattering of GFP fluorescence by the lipid-dense white matter of the 300 µm-thick slices.

To visualise dendritic arborisation, after recording cells were held at their resting membrane potential (RMP) and filled with biocytin-containing internal solution for 30 minutes. Slices with filled neurons were then fixed in 4% paraformaldehyde (PFA) at 4°C overnight. Slices were blocked and permeabilised with a solution of 1% (w/v) bovine serum albumin (BSA) and 0.3% (v/v) Triton X-100 for one hour. They were subsequently incubated with a Streptavidin-Alexa Fluor 594 conjugate (Thermo Fisher Scientific), and anti-GFP Alexa Fluor 488 conjugate antibody (Thermo Fisher Scientific), both diluted 1:1000 in PBS, overnight at 4°C. Semi-assisted traces of the dendrites of filled neurons were made with the ImageJ plugin SNT^18^, which was used to measure and calculate dendritic parameters and perform Sholl analysis.

Input resistance (*R_in_*) was calculated by measuring membrane potential (*V_m_*) near the end of the two hyperpolarizing current steps (*I_cmd_*); the slope of the *V_m_ vs I_cmd_* plot was then calculated to yield *R_in_*. RMP was defined as the mean *V_m_* measured over a 50 ms window immediately before the start of the current step. Rheobase was defined as the amplitude of the smallest 1 s-long current step that elicited at least one action potential (AP). Single AP properties were measured for the first AP that occurred at least 10 ms after the beginning of a current step immediately above rheobase. AP voltage threshold was defined as the *V_m_* at which *dV_m_/dt* first exceeded 15 V/s, AP peak was the voltage reached at the peak of the AP, AP halfwidth was defined as the width of the AP halfway between the AP voltage threshold and the AP peak, AP rise time was the time from 10% to 90% of the AP amplitude, and AP decay time was the time from 100% to 50% of the AP amplitude. The properties of the afterhyperpolarization (AHP) were calculated for the AHPs that followed the APs that fired 10 ms after the beginning of the current step or after the burst at one step above rheobase and averaged the collected values. AHP amplitude was calculated as the voltage at the AHP peak minus the voltage threshold of the preceding AP. AHP halfwidth was calculated at half the AHP amplitude. Burst index was calculated by dividing the seventh inter-stimulus interval (ISI) by the first ISI for the current step which produced at least 8 APs during the current step. Average AP frequency *vs* current plots (f-*I* plots) were produced for all current steps.

### 2.8 Statistical Analysis

All statistical analyses were performed using GraphPad Prism (version 8.0.2, GraphPad Software, USA). Data are reported and plotted as mean ± standard error of the mean (SEM), unless otherwise stated. Statistical significance was determined using a two-tailed unpaired Student’s t-test or parametric one-way ANOVA for normally distributed data, or a Mann-Whitney U-test for unpaired data where the Shapiro-Wilk normality test returned a result of P < 0.05. For multiple comparisons, a two-way ANOVA was used. Results were considered statistically significant where P < 0.05. A summary of statistical analyses can be found in Table S2.

## 3. RESULTS

To generate mosaic *Slc35a2* knockout (KO) mice, foetal mice were injected with the pCas-Guide-Slc35a2 and pCAG-GFP plasmids and electroporated at E15±0.5 (Figure 1B). This methodology results in the transfection of progenitors that give rise to pyramidal neurons in layer 2/3 of the cortex^22^. Confocal microscopy of cryosectioned brain slices from electroporated mosaic *Slc35a2* KO mice revealed GFP in cortical layer 2/3 neurons in a subset of mice, illustrating successful transfection for controls and mice co-injected with both plasmids. Sequence analysis of FACS-sorted GFP^+^ cells collected from neonatal control and mosaic *Slc35a2* KO mice confirmed 8.8% of reads encoded Cas9-induced indels around the *Slc35a2* guide cut site in mosaic mice. No *Slc35a2* indels or mutations were observed in control mice transfected with only pCAG-GFP, confirming Cas9-mediated disruption of the gene (Table S3). In adult control mice, neurons expressing GFP were only observed in layer 2/3 of the cortex (Figure 1C). However, heterotopic neurons in adult mosaic *Slc35a2* KO mice were present in the deeper layers of the cortex, along the grey-white matter boundary and in the white matter itself (Figure 1D-E). Quantifying the positioning of these neurons revealed that mosaic *Slc35a2* KO mice displayed a significantly reduced proportion of GFP^+^ cells correctly positioned in layer 2/3 (P < 0.001, Figure 1F). Areas of increased Olig2-positive (Olig2^+^) cell density are a key histopathological hallmark of MOGHE^1, 2^. Cells immunoreactive to an Olig2 antibody were quantified in the white matter of control and mosaic *Slc35a2* KO mice, however no difference in Olig2^+^ cell density was observed between the two groups (Figure 1G-H).

Heterotopic neurons in both the white matter and deep layers of the cortex were observed to have variable morphology, as revealed by confocal imaging of their dendritic GFP fluorescence (Figure S1A). In addition, somatic GFP fluorescence was used to measure the size of the somata of correctly positioned transfected neurons in layer 2/3 of the cortex; heterotopic neurons were larger than layer 2/3 control neurons (P < 0.05), although soma size of layer 2/3 neurons did not differ between mosaic *Slc35a2* KO mice and GFP-only controls (Figure S1B). GFP-expressing heterotopic neurons were immunoreactive to an antibody directed against SATB2, a transcription factor expressed by cortical neurons that is involved in cortical development and neuronal maturation^23^. Therefore, SATB2 immunoreactivity was utilised to identify a marker of a subset of mature cortical neurons and demonstrate that these heterotopic neurons are developmentally normal and became misplaced during their migration (Figure S1C). Interestingly, a subset of heterotopic neurons was found to display axonal projections into the cortex from the white matter, and therefore could be positioned to contribute to increased excitability (Figure S1D).

Whole-cell current-clamp recordings were made from GFP^+^ neurons correctly located within the layers 2/3. Low numbers and the difficulty in visualising neurons in the white matter precluded recordings from heterotopic neurons. Neurons from which electrophysiological recordings were taken were later visualised through streptavidin staining to verify that recorded neurons were positive for GFP (Figure 2A). The difference in excitability between neurons from GFP-only control and mosaic *Slc35a2* KO mice was assessed by measuring their action potential (AP) firing responses to depolarising current injections. Overall, neurons from mosaic *Slc35a2* KO mice fired fewer APs at a given injected current compared with control neurons (P < 0.0001, Figure 2B-C). Further, the afterhyperpolarisation (AHP) halfwidth was significantly longer in neurons from mosaic *Slc35a2* KO mice, and burst index (a measure of burst-firing) was significantly reduced (P < 0.01, Figure 2F-G). AP parameters such as peak amplitude, halfwidth, rise time and decay time were also measured but were not significantly different between neurons from control and mosaic *Slc35a2* KO mice (Figure S2A-F). No change in input resistance or AHP amplitude was observed (Figure S2G-H).

**Figure 2.**
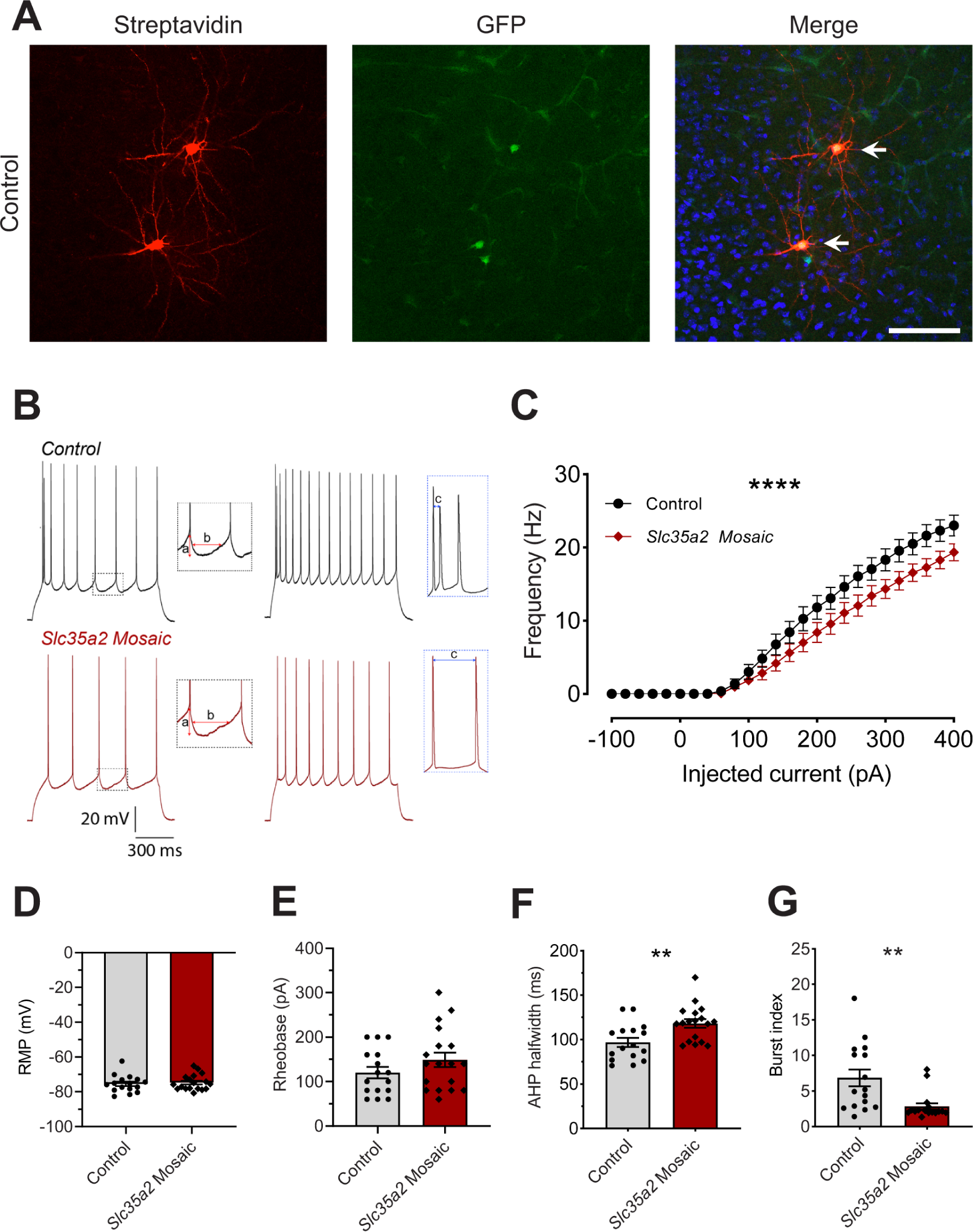
Firing properties of pyramidal neurons are significantly different in mosaic *Slc35a2* KO mice. **(A)** Confocal images of biocytin-filled neurons from a control mouse, showing Streptavidin (red fluorescence) and GFP fluorescence (green), co-stained with DAPI (blue). White arrows indicate recorded neurons positive for both GFP and Streptavidin fluorescence. Scale bar = 100 µm. **(B, top panel)** Example action potentials (APs) elicited by a 140 pA current step (left, black) and by a 200 pA current step (right, black) in pyramidal cells in control mice. **(B, bottom panel)** Example APs elicited by a 220 pA current step (left, red) and by a 280 pA current step (right, red) in pyramidal cells in mosaic *Slc35a2* KO mice. Left inset shows, expanded, the afterhyperpolarisation (AHP) following an AP indicating the parameters measured for analysis: **a** = amplitude of AHP, **b** = halfwidth of AHP. Right inset shows, expanded, initial AP bursts. **c** = time between the first and second APs. **(C)** Averaged action potential frequency-current (*I*) plots in control (n = 16 cells) and in mosaic *Slc35a2* KO (n = 18 cells) mice. Symbols with error bars represent mean ± SEM. **(D-G)** Calculated parameters for membrane potential (RMP), rheobase, AHP halfwidth, and burst index, respectively, in control and mosaic *Slc35a2* KO mice. **P < 0.01, ****P < 0.0001.

Streptavidin staining of filled neurons was subsequently performed to visualise their morphology. Neurons from mosaic *Slc35a2* KO mice showed altered dendritic arborisation compared to control neurons (Figure 3A-B). Sholl analysis of these neurons revealed an increased overall area under the curve and a greater peak of maximal Sholl crossings, suggesting increased dendritic complexity in mosaic neurons (Figure 3C). Additional dendritic parameters significantly increased in mosaic *Slc35a2* KO neurons included total dendrite length (P < 0.05, Figure 3D), number of branches (P < 0.05, Figure 5E), and number of primary (P < 0.01, Figure 3G) and terminal (P < 0.05, Figure 3H) dendritic processes.

**Figure 3.**
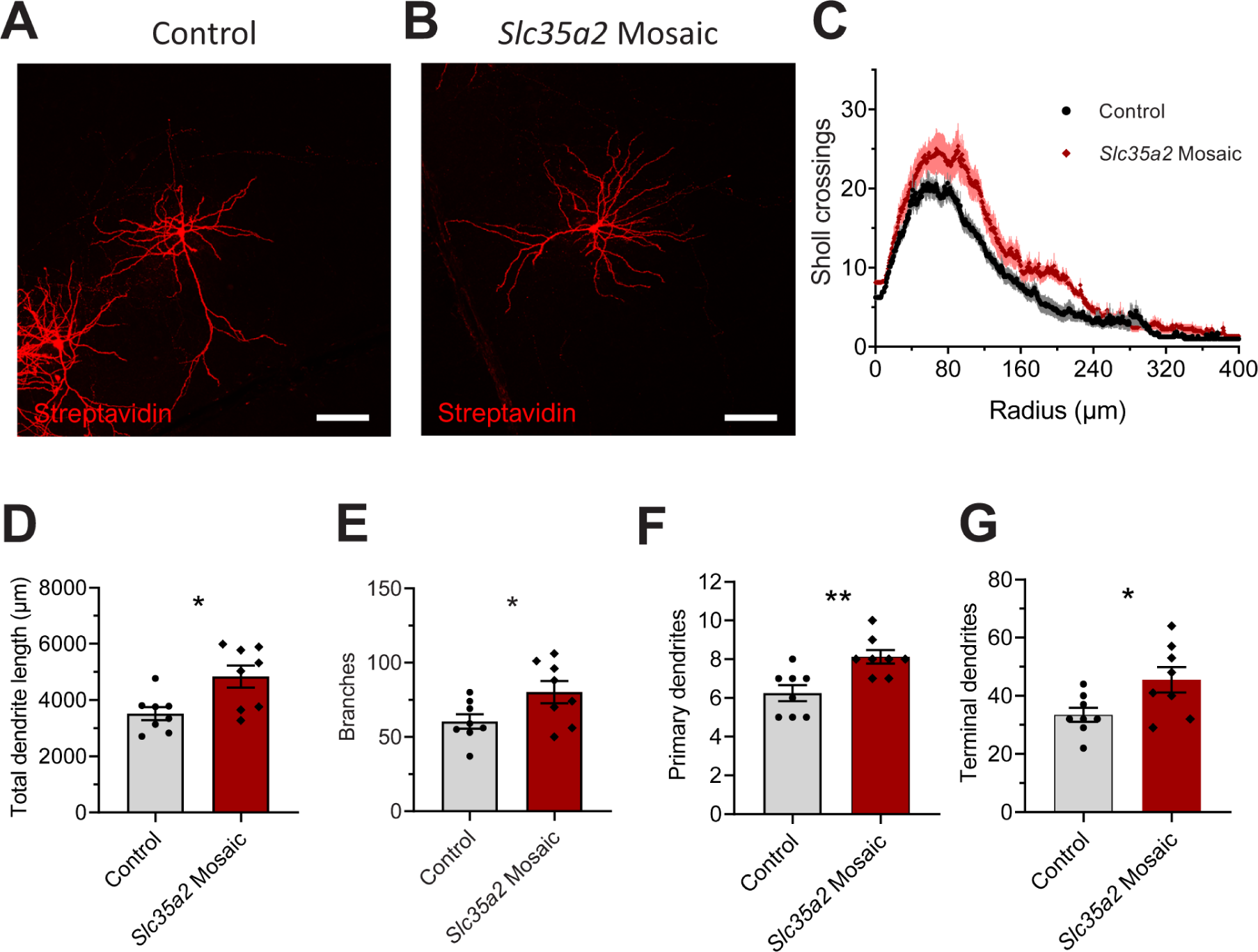
Pyramidal neurons in mosaic *Slc35a2* KO mice display significantly altered dendritic arbor. **(A, B)** Representative confocal images of Biocytin-filled layer 2/3 neurons in control **(A)** and mosaic *Slc35a2* KO **(B)** mice, stained with Streptavidin-594 conjugate. **(C)** Plot of Sholl analyses of dendrite branching in GFP-expressing neurons from control (black) and mosaic *Slc35a2* KO (red) mice. Error bars = SD. **(D-G)** Dendritic parameters measured from neuron traces in control and mosaic *Slc35a2* KO mice. N = 8 cells/group; *P < 0.05, **P < 0.01, **^#^** P < 0.07. Scale bars = 100 µm.

Electrocorticography (ECoG) recordings from mosaic *Slc35a2* KO mice showed frequent large epileptiform spikes with a distinctive waveform (Figure 4A). While a small number of spikes were observed in recordings from GFP-only control mice, both Swiss and C57BL/6J strain mosaic *Slc35a2* KO mice displayed a significantly higher average spike frequency compared with their respective strain-matched controls (P < 0.01 (Swiss), P < 0.05 (C57BL/6J), Figure 4B-C). Spike events in mosaic *Slc35a2* KO mice were not always simultaneous across both hemispheres, with spikes within a single event occurring earlier in one hemisphere suggesting a focal origin (Figure 4D).

**Figure 4.**
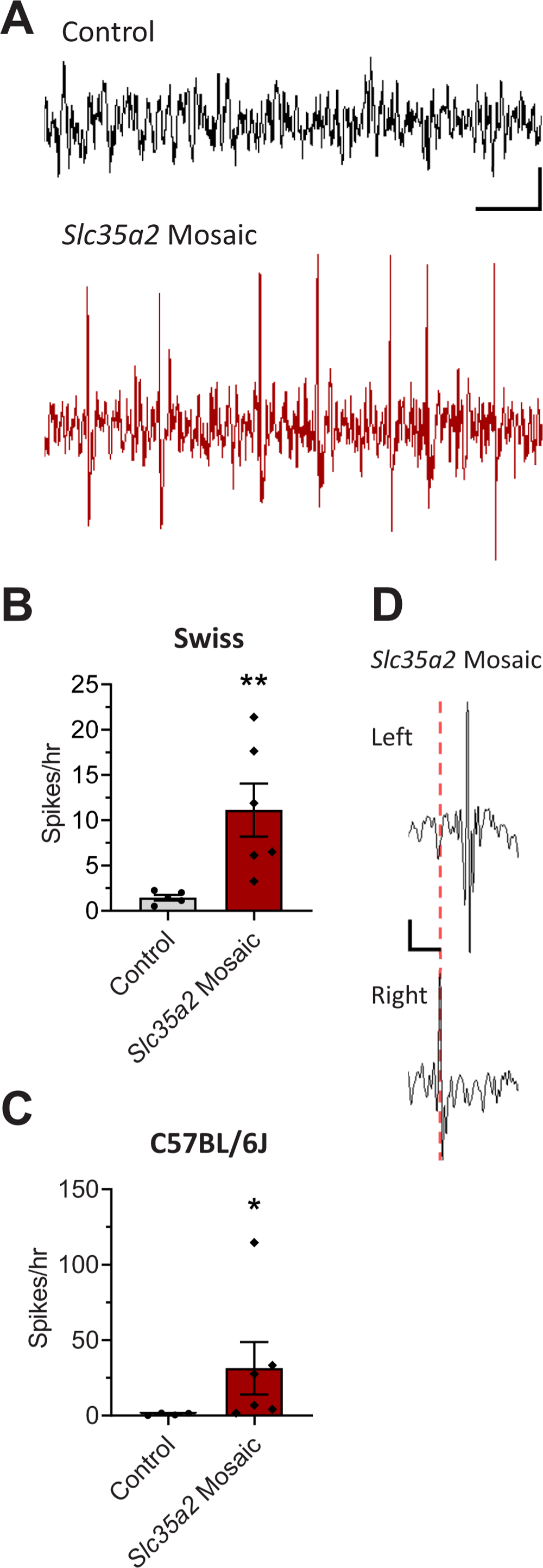
Mosaic *Slc35a2* KO mice display significantly increased epileptiform spike frequencies. **(A)** Sample electrocorticography (ECoG) traces from a control (above, black) and a mosaic *Slc35a2* KO (below, red) mouse, showing examples of bursts of epileptiform spike activity in the latter (scale bar = 2 sec, 100 µV). **(B, C)** Spikes were seen in both Swiss **(B)** and C57BL/6J strain mice **(C)** at a significantly higher frequency in ECoG traces from *Slc35a2* mosiac mice and were very rare in control mice. N = 5 control, 6 mosaic *Slc35a2* KO (Swiss), N = 4 control, 5 mosaic *Slc35a2* KO (C57BL/6J); *P < 0.05, **P < 0.01. **(D)** Expanded view of a spike event in an mosaic *Slc35a2* KO mouse, with traces from the left hemisphere (top) and the right hemisphere (bottom), showing unsynchronised spikes originating in the right hemisphere (scale bar = 200 ms, 100 µV).

Furthermore, mosaic *Slc35a2* KO mice displayed behavioural changes in the open field test. Compared with controls, these mice had significantly increased numbers of ambulatory episodes (P < 0.05, Figure 5A), and demonstrated increased average speed (P < 0.01, Figure 5C) and total distance travelled (P < 0.05, Figure 5D), suggesting a hyperactive phenotype^24^. They also displayed significantly increased time spent in the centre (P < 0.001, Figure 5E), distance travelled in the centre (P < 0.0001, Figure 5F), and entries into the centre zone (P < 0.001, Figure 5H), potentially indicating a decreased anxiety response^25^.

**Figure 5.**
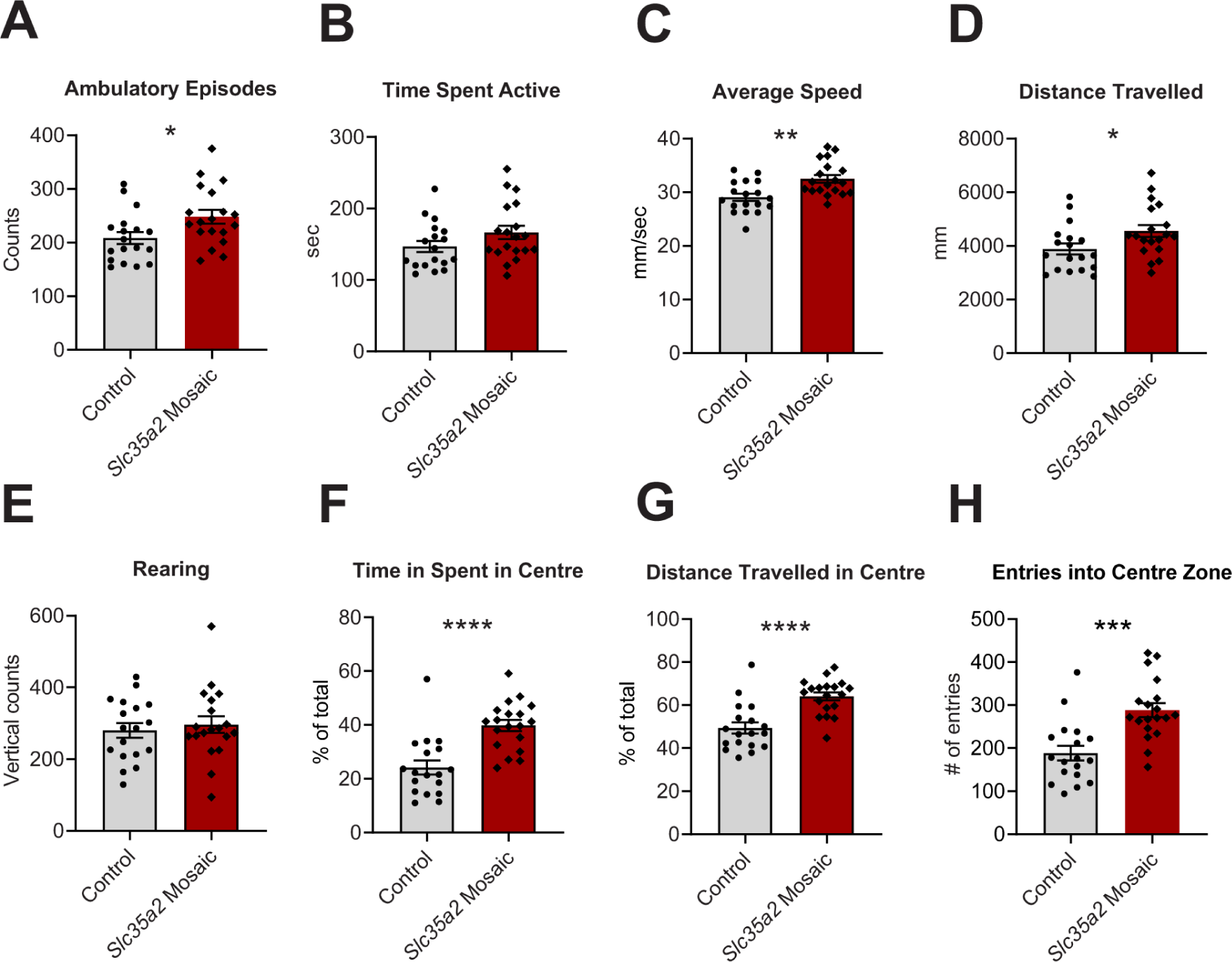
Mosaic *Slc35a2* KO mice display a hyperactive behavioural phenotype. **(A-H)** Open field test locomotion parameters measured in Control and mosaic *Slc35a2* KO mice. Significant increases in speed and ambulation are suggestive of hyperactivity **(A, C, D)**, while significant increases in time and movement in the centre zone of the locomotor cell may suggest a decrease in baseline anxiety **(F-H)**. *P < 0.05. **P < 0.01. ***P < 0.001. ****P < 0.0001.

## 4. DISCUSSION

MOGHE and other focal MCDs such as focal cortical dysplasia (FCD) are a challenging group of disorders to treat, highlighting a need to develop model systems to better understand mechanisms and to act as preclinical tools to identify and test therapeutic interventions. Somatic mosaicism is a common cause of focal MCD with multiple genes now implicated^26^. MOGHE is a relatively recently described MCD with significant clinical sequelae, yet the mechanisms underlying the phenotype remain unclear, especially how a disorder with prominent oligodendroglial abnormalities results in a drug resistant epilepsy. Here we used *in utero* electroporation in combination with CRISPR-mediated mosaic knockout of *Slc35a2* to generate a model that recapitulates some of the key features of MOGHE and provides insights into potential pathological mechanisms.

Brain-specific CRISPR-mediated mosaic knockout of *Slc35a2* during foetal mouse development results in the presence of heterotopic neurons in the white matter. The presence of misplaced neurons in mosaic *Slc35a2* KO mice illustrates that loss-of-function of *Slc35a2* can perturb neuronal progenitor migration during neurodevelopment and produce structural and functional changes. In agreement, similar findings were recently observed using both CRISPR-mediated mosaic knockout of *Slc35a2* and a shRNA knockdown approach^11^. The observation that transient knockdown of *Slc35a2* expression by shRNA results in the same phenotypic outcome as *Slc35a2* KO highlights that downregulation of *Slc35a2* at a critical timepoint during neurodevelopment is sufficient to recapitulate this histopathological aspect of MOGHE^11^. While mosaic *Slc35a2* KO mice did not exhibit spontaneous seizures, they did display a significantly higher frequencies of epileptiform spikes. Similarly, Elziny *et al*. found that mosaic knockdown of *Slc35a2* resulted in an increased sensitivity to proconvulsant seizures^11^. Additionally, mosaic *Slc35a2* KO mice displayed increased locomotion that suggestive of hyperactivity, which is reported at higher rates in other MCDs^27^. While increased density of Olig2^+^ cells is a consistent hallmark in patients with MOGHE^2^, no difference in Olig2^+^ cell density was observed in mosaic *Slc35a2* KO mice or shRNA knock-down^11^. Notably, the IUE protocol utilised in both of these studies specifically targets layer 2/3 neuronal progenitors. This suggests that oligodendroglial density and neuronal progenitor migration perturbations observed in the brain of individuals with MOGHE are likely caused by different, presumably cell-autonomous, mechanisms. Although principal neuron mosaic *Slc35a2* KO alone is sufficient to cause increase neuronal network excitability and behavioural changes, we cannot rule out a role of oligodendrocytes in the pathogenesis of MOGHE.

Biophysical analyses of transfected neurons in the mPFC of mosaic *Slc35a2* KO mice demonstrated that CRISPR-mediated knockdown reduced AP firing frequencies in neurons injected with depolarising currents. While seemingly inconsistent with the increased network-level excitability exhibited by these mice, reduced cortical pyramidal neuron firing has previously been linked to genetic epilepsy. This includes reduced AP firing of dysmorphic neurons from an electroporation-based mouse models of FCD type II that has a robust spontaneous seizure phenotype^14, 15^. It is also well established for some ‘channelopathies’ that loss of function pathogenic variants can cause epilepsy with reduced AP firing in individual neurons occurring concomitantly with increased network-level excitability in model systems^28–30^.

Cortical pyramidal neurons in mosaic *Slc35a2* KO mice showed a significantly longer mean AHP halfwidth, reflecting their decreased intrinsic excitability. In addition, burst-firing was reduced and this may impact micro-network properties^31^. In this capacity, channels responsible for regulating the membrane permeability of Na^+^, K^+^, and Ca^2+^ can have their trafficking, gating and conductance properties impacted by glycosylation and sialylation^9^. The lack of galactose residues on glycan chains in *Slc35a2*-deficient neurons may specifically impair channels that underpin their AHP and burst-firing properties.

The observation of heterotopic neurons in mosaic *Slc35a2* KO mice suggests a key role for this protein in the migration of neuronal progenitors. The lack of UDP-galactose transporter activity would prevent galactose transport into the Golgi, leading to insufficient or aberrant glycosylation of proteins and sphingolipids^5^. As the presence of galactose residues on glycan chains is essential for the linking of sialic acid, which has been shown to play a fundamental role in neural progenitor migration^32^, the disruption of sialylation may underpin the histopathology of MOGHE and the heterotopic neurons observed in mosaic *Slc35a2* KO mice. Similarly, the cytoskeletal regulator MAP2 is known to be glycosylated and incorporate galactose in the central nervous system^33^, and is involved in both dendritic development and cortical migration in association with neural cell adhesion molecule 2 (NCAM2)^34^. NCAM2 is a heavily glycosylated protein, plausibly underpinning an axis by which loss of *Slc35a2* function impacts the neuronal dendritic arbor. The perturbations in cell migration, dendritic arborisation, and neuronal firing may all be impacted by *Slc35a2* deficiency by different mechanisms, which interact to produce the resultant network-level hyperactivity seen in both mosaic *Slc35a2* KO mice and in patients with MOGHE.

All electrophysiology data were recorded from correctly positioned layer 2/3 cortical neurons in P21-25 mice and, in conjunction with the robust dendritic characteristics, strongly support cell-autonomous effects of *Slc35a2* knockout. A limitation of this work is that we did not assess KO efficiency at this timepoint due to the difficulty in isolating intact transfected neurons from mature brain for GFP-mediated FACS isolation. However, validation of Cas9-mediated KO of *Slc35a2* in neonatal mice (P0-2) demonstrated ∼9% incidence of indels detected by sequencing of FACS-enriched cell populations. P21-25 mice are likely to have significantly higher frequency of *Slc35a2* KO in GFP^+^ cells because of the time needed for maximal plasmid expression in electroporated cells^35, 36^, the duration of this expression, and time required for synthesis of the Cas9, translocation to the nucleus and subsequent indel introduction^37, 38^. Furthermore, the role of heterochromatin of the inactive X chromosome in female may also slow Cas9-mediated cutting^30, 31^, with the additional two weeks before analysis at P21-25 providing opportunity for continued indel introduction in transfected cells.

Importantly, *SLC35A2* loss-of-function variants have been detected in both correctly positioned neurons and heterotopic neurons in tissue from MOGHE patients^2^. Therefore, our data suggest that correctly positioned layer 2/3 neurons carrying a variant are likely to contribute to disease. *Slc35a2* KO neurons in layer 2/3 displayed altered dendritic arborisation, a feature that has been associated with increased excitability in other genetic epilepsies^21^. As dendritic arborisation has not yet been assessed in tissue surgically resected from patients with MOGHE, investigation into the morphological and electrophysiological properties of these neurons is warranted.

In summary, we report the generation of mosaic *Slc35a2* KO mice through the utilisation of *in utero* electroporation in combination with CRISPR/Cas9. These mice displayed heterotopic neurons, recapitulating a histopathological hallmark of MOGHE. CRISPR-mediated knockdown of *Slc35a2* in these mice is sufficient to result in increased network excitability in the form of frequent epileptiform spikes as observed on ECoG recordings, and an altered electrophysiological profile characterised by reduced neuronal firing, even in the absence of an observed oligodendroglial perturbation. This suggests the neuronal abnormality seen in MOGHE may be the epileptic driver. At the whole animal level, we observed a hyperactive behavioural phenotype, and increased dendritic arborisation of transfected neurons. In this capacity, our mouse model of *Slc35a2* mosaicism provides a unique opportunity to further investigate the biochemical and biophysical basis of this disease, potentially highlighting precision-based therapeutic strategies.

## Acknowledgements

This work was supported by a National Health and Medical Research Council (NHMRC) Program Grant (10915693) to CR. JS acknowledges the support of an Australian Government Research Training Program Scholarship. HV acknowledges an Al and Val Rosenstrauss Fellowship from the Rebecca L Cooper Foundation. KBH acknowledges support from the Melbourne Children’s Clinician Scientist Fellowship scheme, and grants from the NHMRC and the Medical Research Futures Fund. The Florey Institute of Neuroscience and Mental Health is supported by Victorian State Government Operational Infrastructure Program and the Independent Research Institute Infrastructure Support Scheme.

## Author Contributions

JS and KPA analysed data and prepared figures. JS and CR prepared the manuscript. JS performed *in vivo* experiments, immunohistochemistry, and imaging. KPA performed and analysed brain slice electrophysiology experiments. JS and HV performed FACS and MiSeq experiments, and HV analysed the MiSeq data. CR, SM, PJL, RJL and KBH contributed to the design and direction of the research. All authors contributed to the editing and approved the submission of this manuscript.

## Data Availability

The datasets used and/or analysed during the current study are available from the corresponding author on reasonable request.

## REFERENCES

1. Schurr J, Coras R, Rossler K, et al. Mild Malformation of Cortical Development with Oligodendroglial Hyperplasia in Frontal Lobe Epilepsy: A New Clinico-Pathological Entity. Brain Pathol. 2017 Jan;27(1):26–35.

2. Bonduelle T, Hartlieb T, Baldassari S, et al. Frequent SLC35A2 brain mosaicism in mild malformation of cortical development with oligodendroglial hyperplasia in epilepsy (MOGHE). Acta Neuropathol Commun. 2021 Jan 6;9(1):3.

3. Garganis K, Gkiatis K, Maletic J, et al. Frontal lobe epilepsy and mild malformation with oligodendroglial hyperplasia: Further observations on electroclinical and imaging phenotypes, and surgical perspectives. Epileptic Disord. 2023 Jun;25(3):343–59.

4. Fleischer B. Mechanism of glycosylation in the Golgi apparatus. J Histochem Cytochem. 1983 Aug;31(8):1033–40.

5. Conte F, van Buuringen N, Voermans NC, Lefeber DJ. Galactose in human metabolism, glycosylation and congenital metabolic diseases: Time for a closer look. Biochim Biophys Acta Gen Subj. 2021 Aug;1865(8):129898.

6. Oelmann S, Stanley P, Gerardy-Schahn R. Point mutations identified in Lec8 Chinese hamster ovary glycosylation mutants that inactivate both the UDP-galactose and CMP-sialic acid transporters. The Journal of biological chemistry. 2001 Jul 13;276(28):26291–300.

7. Schwarz F, Aebi M. Mechanisms and principles of N-linked protein glycosylation. Curr Opin Struct Biol. 2011 Oct;21(5):576–82.

8. Scott H, Panin VM. The role of protein N-glycosylation in neural transmission. Glycobiology. 2014 May;24(5):407–17.

9. Scott H, Panin VM. N-glycosylation in regulation of the nervous system. Adv Neurobiol. 2014;9:367–94.

10. Schnaar RL, Gerardy-Schahn R, Hildebrandt H. Sialic acids in the brain: gangliosides and polysialic acid in nervous system development, stability, disease, and regeneration. Physiol Rev. 2014 Apr;94(2):461–518.

11. Elziny S, Sran S, Yoon H, et al. Loss of Slc35a2 alters development of the mouse cerebral cortex. bioRxiv. 2023:2023.11.29.569243.

12. Mantegazza M, Auvin S, Barker-Haliski M, et al. A companion to the preclinical common data elements for rodent genetic epilepsy models. A report of the TASK3-WG1B: Paediatric and genetic models working group of the ILAE/AES joint translational TASK force. Epilepsia Open. 2022 Aug 11.

13. Kilkenny C, Browne W, Cuthill IC, Emerson M, Altman DG, Group NCRRGW. Animal research: reporting in vivo experiments: the ARRIVE guidelines. Br J Pharmacol. 2010 Aug;160(7):1577–9.

14. Hsieh LS, Wen JH, Claycomb K, et al. Convulsive seizures from experimental focal cortical dysplasia occur independently of cell misplacement. Nat Commun. 2016 Jun 1;7:11753.

15. Hsieh LS, Wen JH, Nguyen LH, et al. Ectopic HCN4 expression drives mTOR-dependent epilepsy in mice. Sci Transl Med. 2020 Nov 18;12(570).

16. Aubrey BJ, Kelly GL, Kueh AJ, et al. An inducible lentiviral guide RNA platform enables the identification of tumor-essential genes and tumor-promoting mutations in vivo. Cell Rep. 2015 Mar 3;10(8):1422–32.

17. Bleakley LE, McKenzie CE, Soh MS, et al. Cation leak underlies neuronal excitability in an HCN1 developmental and epileptic encephalopathy. Brain. 2021 Aug 17;144(7):2060–73.

18. Arshadi C, Gunther U, Eddison M, Harrington KIS, Ferreira TA. SNT: a unifying toolbox for quantification of neuronal anatomy. Nat Methods. 2021 Apr;18(4):374–7.

19. Nguyen LH, Xu Y, Nair M, Bordey A. The mTOR pathway genes mTOR, Rheb, Depdc5, Pten, and Tsc1 have convergent and divergent impacts on cortical neuron development and function. bioRxiv. 2023:2023.08.11.553034.

20. Pinares-Garcia P, Spyrou J, McKenzie CE, et al. Antidepressant-like activity of a brain penetrant HCN channel inhibitor in mice. Front Pharmacol. 2023;14:1159527.

21. Reid CA, Leaw B, Richards KL, et al. Reduced dendritic arborization and hyperexcitability of pyramidal neurons in a Scn1b-based model of Dravet syndrome. Brain. 2014 Jun;137(Pt 6):1701–15.

22. Molyneaux BJ, Arlotta P, Menezes JR, Macklis JD. Neuronal subtype specification in the cerebral cortex. Nat Rev Neurosci. 2007 Jun;8(6):427–37.

23. Britanova O, de Juan Romero C, Cheung A, et al. Satb2 is a postmitotic determinant for upper-layer neuron specification in the neocortex. Neuron. 2008 Feb 7;57(3):378–92.

24. Umezu T, Kita T, Morita M. Hyperactive behavioral phenotypes and an altered brain monoaminergic state in male offspring mice with perinatal hypothyroidism. Toxicol Rep. 2019;6:1031–9.

25. Simon P, Dupuis R, Costentin J. Thigmotaxis as an index of anxiety in mice. Influence of dopaminergic transmissions. Behav Brain Res. 1994 Mar 31;61(1):59–64.

26. McConnell MJ, Moran JV, Abyzov A, et al. Intersection of diverse neuronal genomes and neuropsychiatric disease: The Brain Somatic Mosaicism Network. Science. 2017 Apr 28;356(6336).

27. Curatolo P, Moavero R, de Vries PJ. Neurological and neuropsychiatric aspects of tuberous sclerosis complex. Lancet Neurol. 2015 Jul;14(7):733–45.

28. Tamura S, Nelson AD, Spratt PWE, et al. CRISPR activation rescues abnormalities in SCN2A haploinsufficiency-associated autism spectrum disorder. bioRxiv. 2022:2022.03.30.486483.

29. Ogiwara I, Miyamoto H, Tatsukawa T, et al. Nav1.2 haplodeficiency in excitatory neurons causes absence-like seizures in mice. Commun Biol. 2018;1:96.

30. Mao M, Mattei C, Rollo B, et al. Distinctive in vitro phenotypes in iPSC-derived neurons from patients with gain- and loss-of-function SCN2A developmental and epileptic encephalopathy. The Journal of Neuroscience. 2023:JN-RM-0692-23.

31. de Kock CP, Sakmann B. High frequency action potential bursts (>or= 100 Hz) in L2/3 and L5B thick tufted neurons in anaesthetized and awake rat primary somatosensory cortex. J Physiol. 2008 Jul 15;586(14):3353–64.

32. Angata K, Huckaby V, Ranscht B, Terskikh A, Marth JD, Fukuda M. Polysialic acid-directed migration and differentiation of neural precursors are essential for mouse brain development. Mol Cell Biol. 2007 Oct;27(19):6659–68.

33. Ding M, Vandre DD. High molecular weight microtubule-associated proteins contain O-linked-N-acetylglucosamine. The Journal of biological chemistry. 1996 May 24;271(21):12555–61.

34. Parcerisas A, Pujadas L, Ortega-Gasco A, et al. NCAM2 Regulates Dendritic and Axonal Differentiation through the Cytoskeletal Proteins MAP2 and 14-3-3. Cereb Cortex. 2020 May 18;30(6):3781–99.

35. Song B, Kang CY, Han JH, Kano M, Konnerth A, Bae S. In vivo genome editing in single mammalian brain neurons through CRISPR-Cas9 and cytosine base editors. Comput Struct Biotechnol J. 2021;19:2477–85.

36. Dean DA, Machado-Aranda D, Blair-Parks K, Yeldandi AV, Young JL. Electroporation as a method for high-level nonviral gene transfer to the lung. Gene Ther. 2003 Sep;10(18):1608–15.

37. Brinkman EK, Chen T, de Haas M, Holland HA, Akhtar W, van Steensel B. Kinetics and Fidelity of the Repair of Cas9-Induced Double-Strand DNA Breaks. Mol Cell. 2018 Jun 7;70(5):801–13 e6.

38. Straub C, Granger AJ, Saulnier JL, Sabatini BL. CRISPR/Cas9-mediated gene knock-down in post-mitotic neurons. PloS one. 2014;9(8):e105584.

