## Supplemental Material for "*Slc35a2* mosaic knockout impacts cortical development, dendritic arborisation, and neuronal firing in the developing brain"

**SUPPLEMENTAL FIGURES AND TABLES**

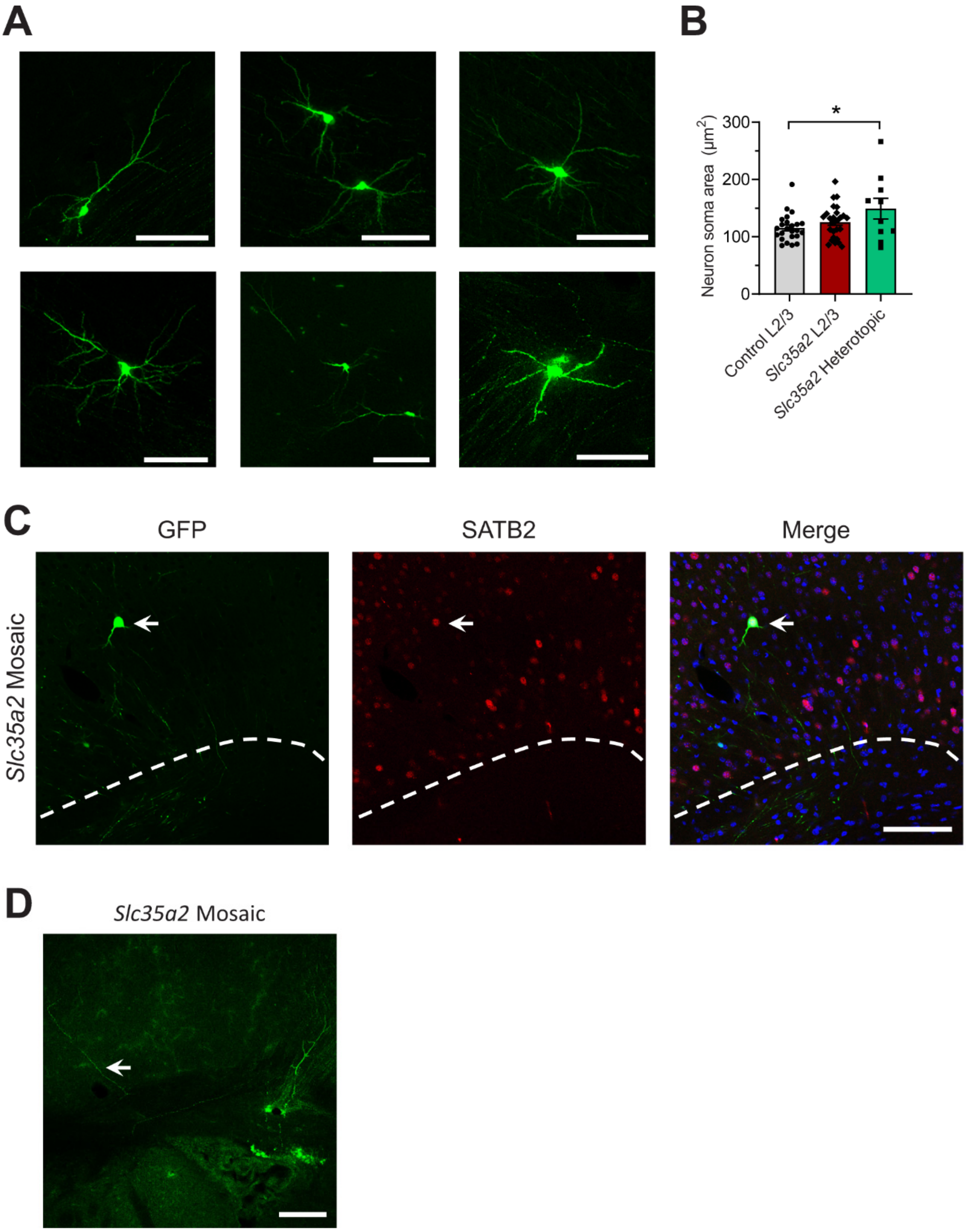


**Figure S1.** **(A)** Representative confocal images showing GFP fluorescence of heterotopic neurons in the white matter and deep layers of the cortex in mosaic *Slc35a2* KO mice. **(B)** Soma area of GFP-expressing neurons in layer 2/3 of the cortex in control and mosaic *Slc35a2* KO mice, and heterotopic neurons in mosaic *Slc35a2* KO mice. *P < 0.05. **(C)** Confocal images for GFP and SATB2 fluorescence in *Slc35a2* mosaic mouse brain tissue co-stained with DAPI, illustrating co-localization of SATB2 immunoreactivity and GFP fluorescence in a heterotopic neuron (white arrow). White dotted line approximates grey-white matter boundary. **(D)** Confocal image of a heterotopic neuron in a mosaic *Slc35a2* KO mouse, showing an axonal projection back into the cortex of the same hemisphere (white arrow). Scale bars = 100 µm.

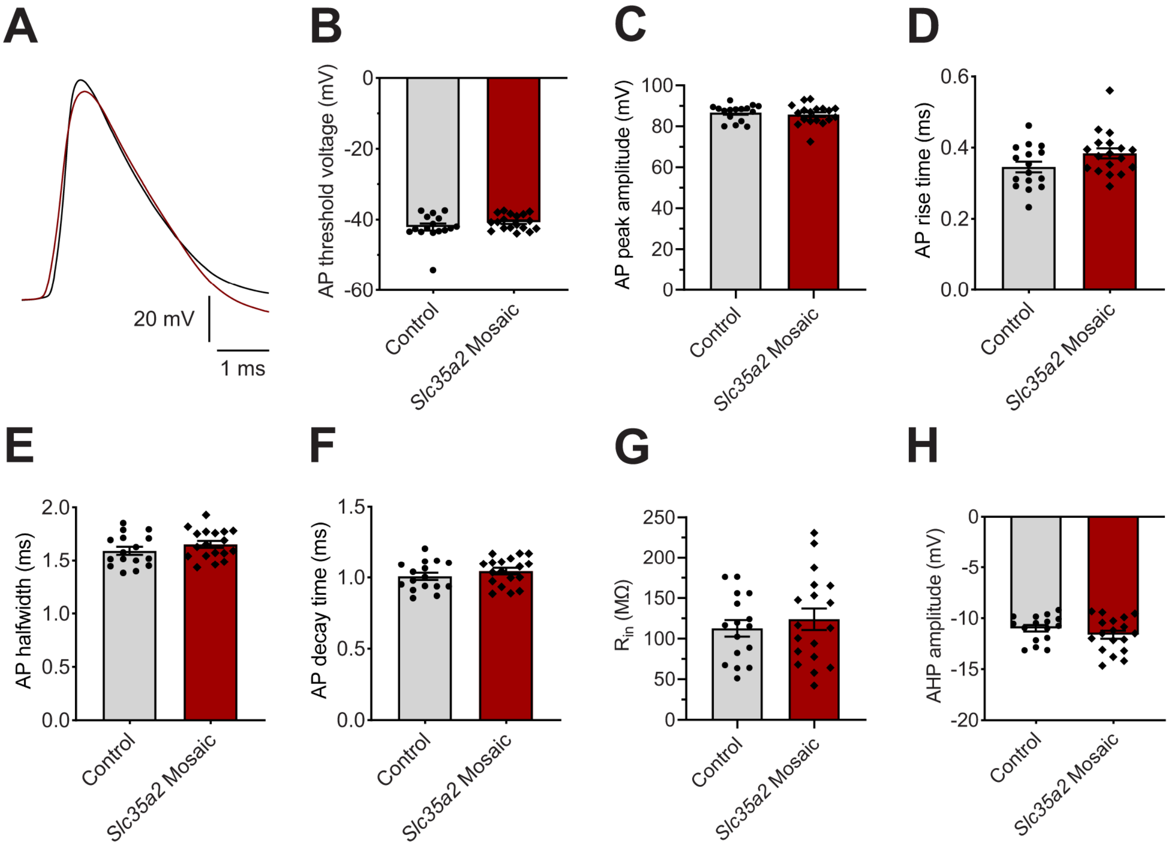


**Figure S2. Action potential (AP) properties do not differ between control and mosaic *Slc35a2* KO mice. (A)** Traces of the first AP recorded from pyramidal neurons, elicited by a 140 pA current step in a control pyramidal neuron (black) and by a 220 pA current step in an *Slc35a2* mosaic KO neuron (red), shown expanded and superimposed. Control AP is the first after the initial burst. **(B-F)** Calculated parameters for AP threshold voltage, AP peak amplitude, AP rise time, AP halfwidth, and AP decay time, respectively, in control (black, n = 16 cells) and in mosaic *Slc35a2* KO (red, n = 18 cells) mice, showing no significant differences. **(D-H)** Similar results for input resistance ($R_{in}$) and afterhyperpolarisation (AHP) amplitude. Data are represented as mean ± SEM.

**Table S1.** Primary antibodies, secondary antibodies, and fluorescent probes used.

| Antibody | Company | Catalog Number | Host Animal | Concentration Used |
|---|---|---|---|---|
| *Primary* | | | | |
| Olig2 | Abcam | ab109186 | Rabbit | 1:100 |
| SATB2 | Abcam | ab51502 | Mouse | 1:125 |
| α GFP, AF488 conjugate | Invitrogen | A-21311 | Rabbit | 1:1000 |
| Streptavidin, AF594 conjugate | Invitrogen | S-32356 | N/A | 1:1000 |
| DAPI | Sigma Aldrich | D9452 | N/A | 1 μg/mL |
| *Secondary* | | | | |
| α Rabbit IgG, AF594 | Invitrogen | A-11012 | Goat | 1:1000 |
| α Mouse IgG1, AF594 | Jackson Immuno | 115-585-205 | Goat | 1:1000 |

**Table S2.** Summary of statistical analysis results. Statistically significant P-values ($P < 0.05$) are indicated by bold type. df = degrees of freedom; t = t-statistic; U = Mann-Whitney U-statistic; F = F-statistic; q = q-statistic; $P_{adj}$ = adjusted P-value. For analyses of variance, the mean squared residual ($MS_{residual}$) is listed under the P column in the Residual row.

[*Spyrou et al Table S2 - Statistical Results.xls*]

**Table S3.** Sequences around the *Slc35a2* cut site, and their proportional abundance, as detected by MiSeq in neonatal (P0-2) control and mosaic *Slc35a2* KO mice. Guide site is shown in blue, additions are shown in red. WT = wild-type sequence; del = deletion; add = addition.

[*Spyrou et al Table S3 - MiSeq Results.xls*]
